# Sex differences in traumatic stress reactivity of rats with a history of alcohol drinking

**DOI:** 10.1101/869990

**Authors:** Lucas Albrechet-Souza, Connor L. Schratz, Nicholas W. Gilpin

## Abstract

**Background:** Alcohol misuse and post-traumatic stress disorder (PTSD) are highly comorbid and treatment outcomes are worse in individuals with both conditions. Although more men report experiencing traumatic events than women, the lifetime prevalence of PTSD is twice as high in females. Despite these data trends in humans, preclinical studies of traumatic stress reactivity have been performed almost exclusively in male animals.

**Methods:** This study was designed to examine sex differences in traumatic stress reactivity in alcohol-naïve rats and rats given intermittent access to 20% ethanol in a 2-bottle choice paradigm for 5 weeks. Rats were exposed to predator odor (bobcat urine) and tested for avoidance of the odor-paired context 24 hours later; unstressed Controls were never exposed to odor. Two days after stress, we measured physiological arousal using the acoustic startle response (ASR) test. We also measured anxiety-like behavior using the elevated plus-maze (EPM) and circulating corticosterone levels before and immediately after odor exposure.

**Results:** Male and female rats exposed to predator odor displayed blunted weight gain 24 hours post-stress, but only a subset of stressed animals exhibited avoidance behavior. Chronic intermittent alcohol drinking increased the proportion of Avoiders in males and predator odor exposure increased ASR in these animals. Predator odor stress reduced ASR in females relative to unstressed females and stressed males, regardless of alcohol drinking history. Bobcat urine exposure did not promote persistent anxiety-like behavior, but alcohol-experienced males exhibited reduced activity in the EPM in comparison to alcohol-experienced females.

Furthermore, predator odor increased circulating corticosterone levels in females relative to males and baseline.

**Conclusions:** We report robust sex differences in behavioral and endocrine responses to bobcat urine exposure in adult Wistar rats. Also, chronic moderate alcohol drinking increased traumatic stress reactivity in males but not females. Our findings emphasize the importance of considering sex as a biological variable in the investigation of traumatic stress effects on physiology and behavior.

## Background

Post-traumatic stress disorder (PTSD) is a chronic psychiatric disease that is seen in some but not all individuals after experiencing a life-threatening traumatic event. Major diagnostic criteria for PTSD include re-experiencing the traumatic event, negative affective state, exaggerated startle responses and persistent avoidance of trauma-related cues [1]. Women are twice as likely to develop PTSD after trauma [2, 3] and women with trauma exposure and/or PTSD exhibit more sensitivity to and less tolerance of negative emotions [4, 5].

Alcohol use disorder (AUD) is one of the most common co-occurring conditions among individuals diagnosed with PTSD [6, 7]. Approximately one-third of individuals with lifetime PTSD also meet criteria for AUD [8]. Some populations, such as military personnel, are at high risk for AUD and PTSD comorbidities. For example, in a study of Iraq and Afghanistan veterans, 63% of those diagnosed with AUD also met criteria for PTSD [9]. Although men have a higher prevalence of AUD than women, and women have a higher prevalence of PTSD than men, any individual with either disorder is more likely to have the other [10]. Individuals with co-occurring AUD and PTSD exhibit a more complex and severe clinical profile than those diagnosed with either disorder alone [8, 11, 12]. Some people with PTSD use alcohol in an attempt to ameliorate debilitating symptoms such as anxiety and hyperarousal [13, 14]. However, there is a complex reciprocal relationship between stress and alcohol: while PTSD symptoms may lead an individual to drink more alcohol, alcohol use may exacerbate PTSD symptoms [15–17].

Exaggerated startle response is considered a hallmark symptom of PTSD [1] and is predictive of disease severity [18]. In a study of military service members referred for psychiatric evaluation for suicide-related concerns, the hyperarousal symptom cluster was the only significant predictor of subsequent suicide attempts [19]. Although previous studies have shown elevated autonomic responses to startling tones in trauma survivors [20–22], other investigations of startle responsivity in patients with PTSD have produced mix results. For example, studies of Gulf War veterans with PTSD found both self-reported [23] and physiologically [24] exaggerated startle compared to non-PTSD veterans. On the other hand, Vietnam veterans with PTSD did not show increased startle [25] unless they were subjected to a stressful environment [26]. More extreme, some studies report that PTSD patients exhibit blunted motor reflex responses to acoustic stimuli [27, 28]. These last two studies used either all female subjects [28] or a majority of females [27], suggesting there may be a sex difference in the presentation of ASR in females.

Preclinical studies using animal models to recapitulate PTSD-like behavioral deficits allow investigation of the biological mechanisms underlying traumatic stress effects, but most preclinical research has been conducted in male animals, potentially neglecting issues specific to female subjects [29, 30]. Predator exposure and predator scent are psychological stressors commonly used as animal models of PTSD. Rodents are exposed to predator odor in a variety of ways, including indirect exposure to a predator (cat), predator urine (bobcat, fox), predator feces or litter, or trimethylthiazoline (TMT) – a synthetic compound isolated from fox feces [31–34]. Although predator stress has been shown to elicit lasting increases in freezing and avoidance behaviors [35, 36], the effects of predator odor stress on ASR are not consistent. For example, male rats exhibit potentiated ASR magnitude *during* exposure to TMT [37], but three exposures to cat odor failed to promote persistent changes on startle reactivity [38]. Prior work from our laboratory showed small increases in ASR at low decibel levels after bobcat urine exposure in alcohol-naïve male rats [39].

The current experiments were designed to test sex differences in traumatic stress reactivity in rats with and without a history of chronic voluntary alcohol consumption exposed to a bobcat urine stress model developed in our laboratory [40]. The urine of carnivorous species contains 2-phenylethylamine, a trace amine produced by the breakdown of the amino acid phenylalanine [41], that activates trace amine-associated receptor 4 (TAAR4) in the rodent olfactory cortex and produces avoidance behavior in rodents [41]. The main hypothesis of this study was that bobcat urine exposure would increase acoustic startle reactivity in male and female rats and that this effect would be exaggerated in rats that exhibit avoidance of trauma-paired cues and in rats with a history of alcohol drinking.

## Methods

### Animals

Eight-week-old male and female Wistar rats (total N = 110) (Charles River, Raleigh, NC) were housed in same-sex pairs in a humidity- and temperature-controlled (22 °C) vivarium on a 12-hours reversed light-dark cycle (lights off at 7 a.m.). Animals had *ad libitum* access to food and water throughout the experiments and were handled daily for 1 week before the initiation of experimental protocols. All behavioral testing occurred in the dark phase. Female rats were freely cycling and assigned to treatment groups without regard to estrous cycle stage. At the end of the experiments, rats were sacrificed by decapitation under isoflurane anesthesia.

### Intermittent access 2-bottle choice alcohol homecage drinking

Paired-housed adult male and female Wistar rats were given access to alcohol and water in three 24-hour sessions per week for 5 weeks prior to stress, as previously described [42]. Briefly, rats were weighed and given access to 1 bottle of 20% v/v ethanol and 1 bottle of water approximately 3 hours after the start of the dark cycle on Mondays, Wednesdays and Fridays. After 24 hours, the alcohol bottle was replaced with a second water bottle that was available for the next 24 hours. Over the weekends, rats had unlimited access to 2 water bottles after the alcohol bottle was removed on Saturday. Bottles were weighed 24 hours after alcohol presentation. The position of the alcohol bottle was alternated across sessions to control for side preferences. Blood was collected from the tail 2 hours after the start of alcohol access during the last drinking session to determine blood alcohol concentration. Blood samples were centrifuged at 1900×g for 14 minutes, after which plasma was collected and immediately analyzed using an Analox AM1 analyzer (Analox Instruments).

### Predator odor stress

Rats were tested in a 5-day conditioned place aversion procedure [40] that began after the acclimatization period in alcohol-naïve rats or 24 hours after the last drinking session in alcohol-experienced rats. On the first day, rats were allowed 5 minutes of free exploration of the apparatus (3-chamber pre-test session), which consisted of three large chambers (36 cm length × 30 cm width × 34 cm height) with different types of floor texture (circles, grid or rod floor) and patterned walls (circles, white or stripes), separated by a small triangular connecting chamber. The apparatus was thoroughly cleaned between animals with Quatricide® PV in water at a concentration of 1:64 (Pharmacal Research Labs, Waterbury, CT). For each rat, the chamber that exhibited the most deviant time score of the three (i.e., highly preferred or highly avoided) was excluded from all future sessions for that rat. On day 2, the rat was allowed 5 minutes to explore the two non-excluded conditioning chambers (pre-exposure session). Rats were assigned to predator odor stress or unstressed Control groups that were counterbalanced for the magnitude of baseline preference for one chamber versus the other (i.e., groups were assigned such that mean pre-existing preference for each of the two chambers was approximately zero for Stress and Control groups). For rats in the Stress group, an unbiased and counterbalanced design was used to determine which chamber (i.e., more preferred or less preferred) would be paired with predator odor for each rat. On day 3, each rat was placed in one of the two chambers with the guillotine door shut without odor for 15 minutes (neutral exposure). On day 4, rats were weighed and placed in the other chamber with the guillotine door shut and a sponge soaked with 3 ml of bobcat urine (*Lynx rufus*; Maine Outdoor Solutions, Hermon, ME) placed under the floor for 15 minutes (odor exposure). Control rats were treated identically to odor-exposed rats, but the sponges did not contain bobcat urine. On day 5, rats were weighed and allowed to explore the two chambers for 5 minutes (post-exposure session). All testing was conducted under indirect dim illumination (one 60W white light facing the wall providing approximately 10 lux in the apparatus) and all sessions were recorded and time spent in each chamber was scored by a treatment-blind observer. Avoidance was quantified as a difference score calculated as time spent in odor-paired chamber on day 5 minus time spent in the same chamber on day 2. Rats that displayed >10-s decrease in time spent in the odor-paired context were classified as Avoiders; all other bobcat urine-exposed rats were classified as Non-Avoiders.

### Acoustic startle response (ASR) test

Two days after exposure to predator odor stress (i.e., day 6 of the protocol), rats were tested for acoustic startle reactivity. Rats were placed in a Plexiglas tube attached to an accelerometer inside a dark, soundproof chamber (SR-Lab, San Diego Instruments, CA) and allowed to acclimate for 5 minutes (75-dB background noise) before the test session, as previously described [39]. This background white noise was present throughout the session. The chamber and Plexiglas tube were cleaned with Quatricide between each animal. Before testing, an S-R calibrator tube was used to calibrate the chambers. The test session consisted of 30 trials with startle stimuli of three different decibel levels: a 750-ms burst of 95 dB, 105 dB or 115 dB white noise was presented 10 times each, separated by a 30-s fixed inter-trial interval. The maximum startle response (Vmax, arbitrary units) was recorded during the first 100 milliseconds of each trial.

### Elevated plus-maze (EPM) test

The EPM test was used to test anxiety-like behavior on day 21 (17 days after exposure to bobcat urine). The EPM was a black Plexiglas apparatus consisting of two closed arms (50 cm × 10 cm × 40 cm) and two open arms (50 cm × 10 cm) attached to metal legs elevating the maze 50 cm above the ground. All testing was conducted under dim illumination (approximately 10 lux in the open arms) during the dark phase of the light-dark cycle. Rats were placed individually in the center of the maze facing a closed arm and allowed 5 minutes of free exploration. Behavior was recorded with a video camera positioned above the maze. The EPM was cleaned thoroughly between subjects using Quatricide. Video scoring was done by an observer blind to the conditions; we measured percent time spent in the open arms ((open/open+closed) × 100) and number of closed and open arm entries. One arm entry was defined as all four paws entering the arm.

### Plasma Corticosterone Levels

A separate cohort of alcohol-naïve male and female rats was individually transferred from the home cage to a clean cage and exposed to bobcat urine for 15 minutes. Bobcat urine (∼3 ml) was added to a sponge that was placed beside the cage. Tail blood was collected before and immediately after exposure to predator odor stress in EDTA-covered tubes and samples were centrifuged at 1900×g for 20 minutes. Plasma was stored at −80 °C and analyzed in duplicate for corticosterone levels using a DetectX ELISA kit (Arbor Assays, Ann Arbor, MI) according to the manufacturer’s instructions.

### Statistical analysis

Data are reported as mean ± SEM, except where otherwise indicated. All statistics were run using Prism 8 (GraphPad, La Jolla, CA). Alcohol drinkers and alcohol-naïve animals were not tested in parallel, therefore, they are not analyzed together. Change in time spent in predator odor-paired chamber, body weight gain, percent time spent in the open arms of the EPM and closed arms entries were analyzed with two-way analysis of variance (ANOVA) – the variables in all cases were sex and stress condition. Measures of Vmax were normalized by body weight in kilograms. Three-way repeated measures ANOVAs were performed – the variables were sex, stress condition and decibel level, followed by two-way ANOVAs to test stress and sex effects on startle reactivity at each decibel level. Alcohol consumption and plasma corticosterone were analyzed with two-way repeated measures ANOVAs – the variables were sex and sessions. Fisher’s exact test was used to analyze the proportion of Avoiders and Non-Avoiders in each sex in alcohol-naïve and alcohol-experienced rats. Student’s unpaired t-tests were used to compare the magnitude of avoidance in male and female Avoiders, and the startle response and percent time spent in the open arms of the EPM between Avoiders and Non-Avoiders in each sex in alcohol-naïve and alcohol-experienced rats. In cases of significant ANOVA effects, *post hoc* comparisons were performed using Tukey’s multiple comparisons test. Values of P < 0.05 were considered statistically significant.

## Results

### Predator odor stress reactivity in alcohol-naïve rats

Independent of sex, Avoiders exhibited significantly greater avoidance of the predator odor-paired chamber at 24 hours post-exposure (F (1, 44) = 78.51, P < 0.0001) relative to Non-Avoiders (Fig. 2a). There was no significant difference in the magnitude of avoidance between male and female Avoiders (Fig. 2a; t = 1.86, P = 0.08), and the proportion of animals that met Avoider criteria was similar in both sexes (Fig. 2b; P > 0.05). Likewise, both male and female rats exposed to predator odor exhibited significantly reduced body weight gain relative to unstressed Controls 24 hours post-stress (day 5 minus day 4) (Fig. 2c; stress effect: F (1, 44) = 6.99, P = 0.01) but not 4 days post-stress (stress effect: F (1, 44) = 1.12, P = 0.29; data not shown).

**Fig. 1.**
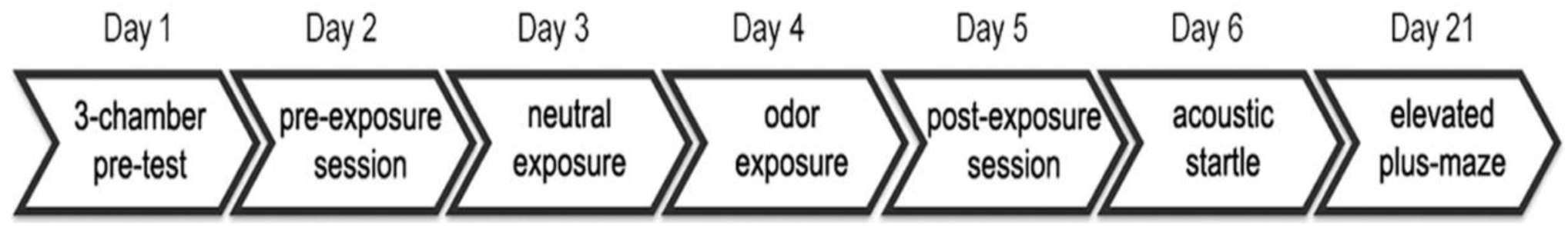
Schematic representation of the experimental design. Male and female rats underwent the conditioned place aversion paradigm using bobcat urine. Avoidance behavior was measured on day 5 (24 hours post-stress). Controls were never exposed to predator odor. Rats with a history of alcohol consumption (intermittent access 2-bottle choice, 5 weeks) were exposed to the same procedure starting 24 hours after the last drinking session. Acoustic startle reactivity was evaluated on day 6 (2 days post-stress) and anxiety-like behavior was tested on day 21 (17 days post-stress).

**Fig. 2.**
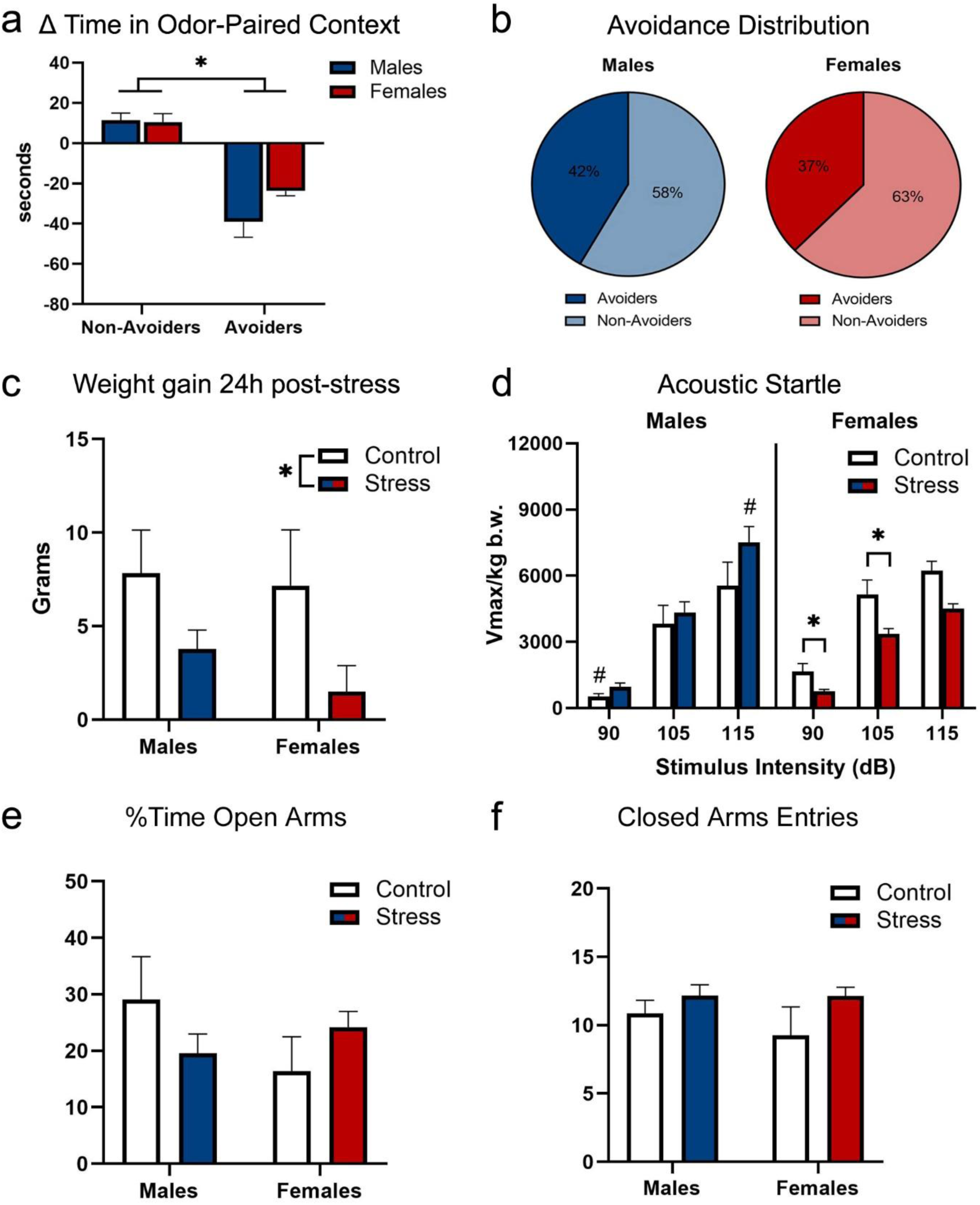
Traumatic stress response in alcohol-naïve rats. **a** Change in time spent in predator odor-paired chamber in male and female rats indexed as Avoiders or Non-Avoiders 24 hours post-stress. **b** Avoidance distribution (Avoiders × Non-Avoiders) in male and female rats. **c** Weight gain in male and female rats measured 24 hours after exposure to predator odor. **d** Acoustic startle response in male and female rats measured 48 hours after exposure to predator odor stress. **e** Percent time spent in the open arms of the elevated plus-maze in male and female rats on day 21 (17 days post-stress). **f** Frequency of entries into the closed arms of the elevated plus-maze in male and female rats on day 21 (17 days post-stress). Data presented as mean ± SEM. * denotes P < 0.05 between indicated groups; # denotes P < 0.05 between sexes.

Because we did not find significant differences in the ASR between Avoider and Non-Avoider males nor between Avoider and Non-Avoider females, these animals were pooled into a single group, designated Stress, and compared to unstressed Controls (Fig. 2d). A three-way repeated measures ANOVA yielded a significant main effect of decibel level (F (2, 116) = 147.50, P < 0.0001), a decibel level × sex interaction effect (F (2, 116) = 4.45, P = 0.01) and a sex × stress interaction effect (F (1, 116) = 7.69, P = 0.01) on ASR. To determine whether predator odor stress affected startle reactivity differently in male and female rats, ASR data for each decibel were analyzed with a two-way ANOVA. At 90 dB, we found a significant main effect of sex (F (1, 58) = 5.45, P = 0.02) and a sex × stress interaction effect (F (1, 58) = 10.59, P = 0.002) on ASR. Tukey’s *post-hoc* comparisons revealed that control females exhibited higher startle reactivity than control males (P = 0.01) and that stressed females showed lower startle reactivity relative to control females (P = 0.01) in response to a 90 dB stimulus. At 105 dB, the two-way ANOVA revealed a significant sex × stress interaction effect (F (1, 58) = 4.11, P = 0.047) on ASR. Tukey’s *post-hoc* comparison revealed that stressed females exhibited lower startle reactivity in response to a 105 dB stimulus than control females (P = 0.02). At 115 dB, the two-way ANOVA revealed a significant sex × stress interaction effect (F (1, 58) = 6.02, P = 0.02) on ASR. Tukey’s *post-hoc* comparison revealed that stressed males exhibited higher startle reactivity in response to a 115 dB stimulus than stressed females (P = 0.0008).

On day 21, rats were tested for anxiety-like behavior in the EPM. Again, we did not find significant differences between Avoider and Non-Avoider males nor between Avoider and Non-Avoider females on percent time spent in the open arms of the EPM; thus, these animals were pooled into a single group, designated Stress, and compared to unstressed Controls. A two-way ANOVA revealed no significant effect of sex (F (1, 59) = 0.73, P > 0.05) or stress (F (1, 59) = 0.04, P > 0.05) on percent time spent in the open arms of the EPM (Fig. 2e). General locomotor performance was assessed by counting closed-arm entries in the EPM. Neither sex (F (1, 59) = 0.59, P > 0.05) nor stress (F (1, 59) = 3.65, P > 0.05) affected number of closed arms entries (Fig. 2f).

### Predator odor stress reactivity in rats with a history of alcohol drinking

Before exposure to predator odor stress, pair-housed rats were given intermittent access to alcohol 20% v/v and water for 5 weeks in the homecage. Because rats were pair-housed, we did not measure intake for individual animals but instead for each cage. We chose this procedure to avoid exposing animals to social isolation stress. A two-way repeated measures ANOVA showed that males consumed significantly higher quantities of alcohol than females over the weeks (Fig 3c; F (1, 15) = 15.49, P = 0.001). However, males and females did not exhibit different blood alcohol levels (BALs) analyzed 2 hours after the beginning of the last alcohol drinking session (t = 0.21, P = 0.83; data not shown). Independent of sex, Avoiders with an alcohol drinking history exhibited significantly greater avoidance of the predator odor-paired chamber at 24 hours post-exposure (F (1, 18) = 28.79, P < 0.0001) relative to Non-Avoiders with an alcohol drinking history (Fig. 3a). There was no significant difference in the magnitude of avoidance between male and female Avoiders (Fig. 3a; t = 0.76, P = 0.48). Although the difference in the proportion of Avoiders and Non-Avoiders in male and female rats did not reach statistical significance (P = 0.39) due to sample size (given a 95% confidence level and 80% power, the recommended sample size would be n = 95), the proportion of alcohol-drinking males that met Avoider criteria was 27% higher than alcohol-drinking females (Fig. 3b) and 25% higher than alcohol-naïve males (compare to data in Fig. 2b).

**Fig. 3.**
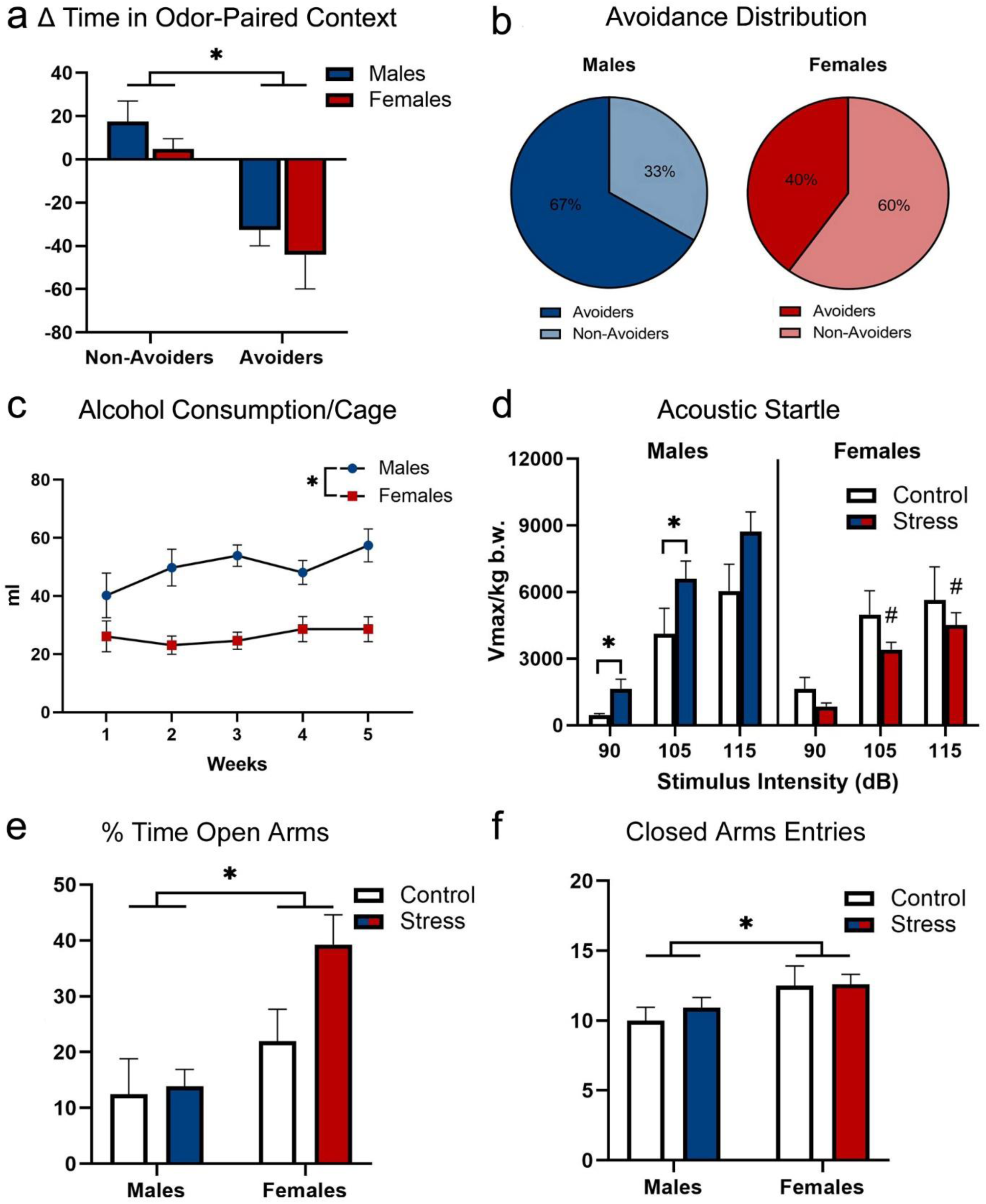
Traumatic stress response in rats given intermittent access to 20% ethanol in a 2-bottle choice paradigm for 5 weeks. **a** Change in time spent in predator odor-paired chamber in male and female rats indexed as Avoiders or Non-avoiders 24 hours post-stress. **b** Avoidance distribution (Avoiders × Non-Avoiders) in male and female rats. **c** Weekly alcohol consumption per cage in same-sex paired-housed male and female rats. **d** Acoustic startle response in male and female rats measured 48 hours after exposure to predator odor stress. **e** Percent time spent in the open arms of the elevated plus-maze in male and female rats on day 21 (17 days post-stress). **f** Frequency of entries into the closed arms of the elevated plus-maze in male and female rats on day 21 (17 days post-stress). Data presented as mean ± SEM. * denotes P < 0.05 between indicated groups; # denotes P < 0.05 between sexes.

In males and females with an alcohol drinking history, Avoider and Non-Avoider did not exhibit differences in ASR, so these animals were pooled into a single group, designated Stress, and compared to unstressed Controls (Fig. 3d). A three-way repeated measures ANOVA yielded a significant main effect of decibel level (F (2, 58) = 130.00, P < 0.0001), a decibel level × sex interaction effect (F (2, 58) = 7.42, P = 0.001) and a sex × stress interaction effect (F (1, 58) = 5.62, P = 0.02) on ASR. To determine whether predator odor stress affected startle reactivity differently in male and female rats with a history of alcohol drinking, ASR data for each decibel were analyzed with a two-way ANOVA. At 90 dB, we found a significant sex × stress interaction effect (F (1, 29) = 7.02, P = 0.01) on ASR. Tukey’s *post-hoc* comparison revealed that stressed males exhibited higher startle reactivity in response to a 90 dB stimulus than control males (P = 0.03). At 105 dB, the two-way ANOVA revealed a significant sex × stress interaction effect (F (1, 29) = 5.94, P = 0.02) on ASR. Tukey’s *post-hoc* comparisons revealed that stressed males exhibited higher startle reactivity in response to a 105 dB stimulus relative to control males (P = 0.04) and also relative to stressed females (P = 0.004). At 115 dB, the two-way ANOVA revealed a significant main effect of sex (F (1, 29) = 5.23, P = 0.03) on ASR. Tukey’s *post-hoc* comparison revealed that stressed males exhibited higher startle reactivity in response to a 115 dB stimulus than stressed females (P = 0.01).

On day 21, rats were tested for anxiety-like behavior in the EPM. We did not find significant differences between Avoider and Non-Avoider males nor between Avoider and Non-Avoider females on percent time spent in the open arms of the EPM; thus, these animals were pooled into a single group, designated Stress, and compared to unstressed Controls. A two-way ANOVA revealed that regardless of stress condition, males with a history of alcohol drinking spent less time in the open arms of the EPM relative to females (Fig. 3e; F (1, 30) = 11.86, P = 0.002), likely driven by the apparent stress-induced increase in time spent in the open arms of the EPM in females. Males with a history of alcohol drinking also exhibited fewer closed arm entries compared to females (Fig. 3f; F (1, 30) = 4.96, P = 0.03).

### Predator odor stress effects on plasma corticosterone

Predator odor stress produced sexually dimorphic changes in circulating corticosterone levels (Fig. 4). A two-way repeated measures ANOVA yielded a significant main effect of sex (F (1, 9) = 17.43, P = 0.002), time (F (1, 9) = 62.40, P < 0.0001) and a sex × time interaction effect (F (1, 9) = 20.91, P = 0.001) on plasma corticosterone measured before and immediately after a 15-min exposure to bobcat urine. Tukey’s *post-hoc* comparison revealed that, after bobcat urine exposure, circulating corticosterone levels were significantly higher in stressed females relative to stressed males (P = 0.0001) and also relative to their own baseline (P = 0.001).

**Fig. 4.**
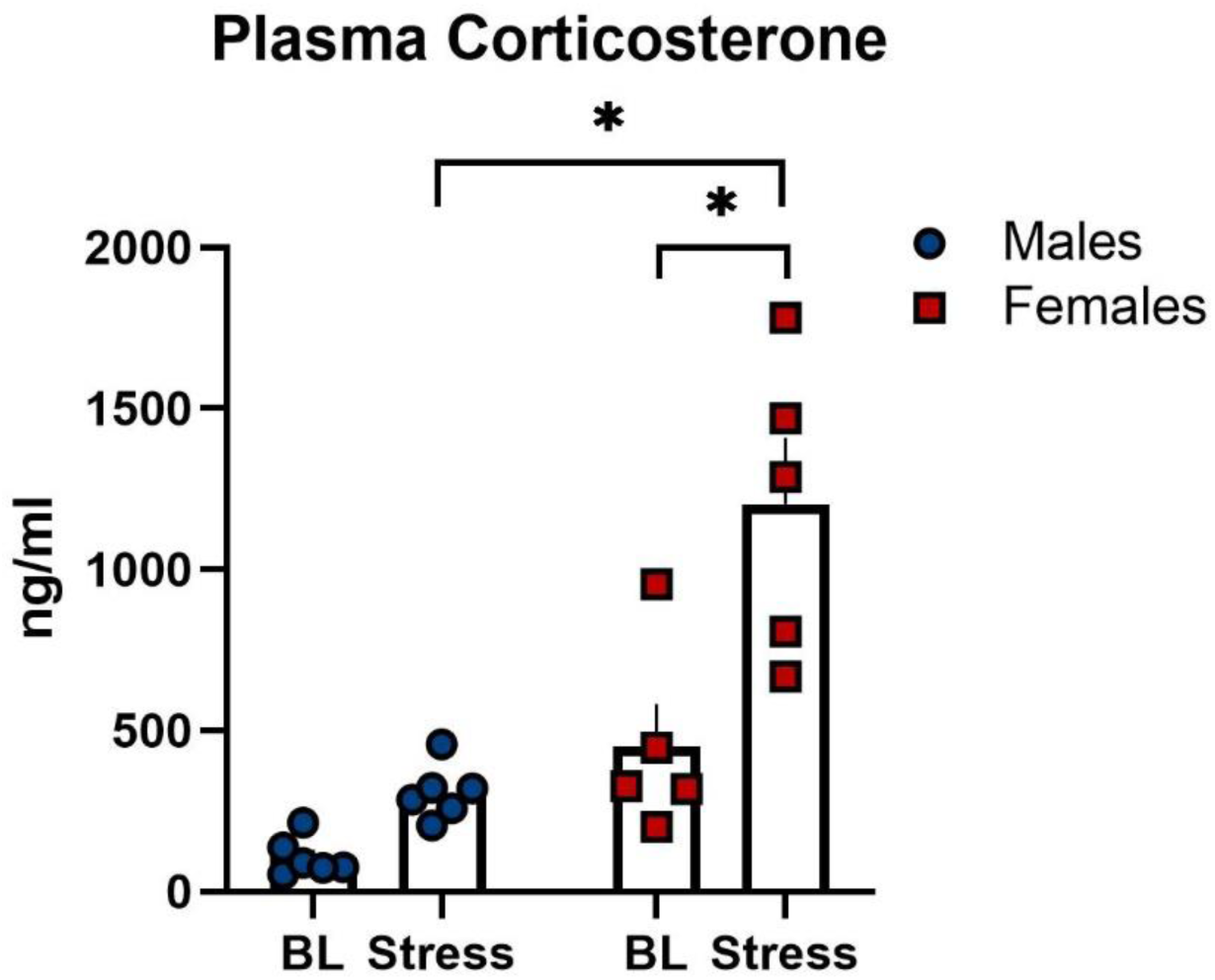
Plasma corticosterone levels measured at baseline (BL) and immediately following 15 minutes exposure to bobcat urine (Stress) in male and female rats. Data presented as mean ± SEM. * denotes P < 0.05.

## Discussion

We report that alcohol-naïve male and female rats exposed to predator odor displayed blunted weight gain 24 hours post-stress even though only a subset of stressed animals exhibited avoidance behavior. A similar percentage of alcohol-naïve males and females were classified as Avoiders after stress, but the proportion of Avoiders was much higher in alcohol-experienced males relative to females with a history of alcohol drinking and alcohol-naïve males. Predator odor exposure enhanced startle reactivity in alcohol-experienced males only, but regardless of alcohol drinking history, stress reduced startle reactivity in females. Predator odor stress did not produce anxiety-like behavior, but alcohol-experienced males spent less time in the open arms than females and exhibited less locomotor activity than females in the EPM. Furthermore, bobcat urine exposure significantly increased circulating corticosterone levels in females relative to males and baseline. These results are important because alcohol use exacerbates PTSD symptoms in humans [15–17] and risk factors for co-morbid PTSD and AUD may differ based on sex [12].

Predator odor stress produces behavioral, molecular, and physiological alterations that recapitulate some PTSD symptoms [40, 43]. In line with previous studies, bobcat urine exposure elicited significant avoidance in a subset of animals [35, 44]. A similar proportion of alcohol-naïve male and female rats exposed to predator odor were classified as Avoiders (males: 42%; females: 37%), with similar magnitude of bobcat urine-paired context avoidance 24 hours post-stress. After chronic voluntary ethanol consumption, a much higher percentage of males (but not females) were classified as Avoiders (males: 67%; females: 40%). A systematic review of the comorbidity between PTSD and alcohol misuse found associations between alcohol consumption and the avoidance/numbing and hyperarousal PTSD symptom clusters [45]. We previously reported that male Avoider rats exhibit persistent increases in alcohol self-administration and that avoidance behavior predicts post-stress escalation of alcohol drinking [35].

Although only a subset of rats exposed to predator odor displayed avoidance behavior, all stressed male and female rats exhibited signs of physiological stress. A previous study from our laboratory demonstrated that all male rats exposed to predator odor (i.e., Avoiders and Non-Avoiders) exhibit anxiety-like behavior measured 2 days and 5 days post-stress [46]. The current data build on that work by reporting that all alcohol-naïve stressed male and female rats displayed blunted weight gain 24 hours after predator odor exposure (we did not evaluate changes in body weight in rats with a history of alcohol drinking). Moreover, prior work from our laboratory showed that male rats exposed to bobcat urine exhibit a non-significant general increase in startle reactivity [39]. In the present study, we confirmed and extended these findings by testing females and animals with a history of chronic voluntary alcohol drinking.

The startle response is an operational measure of threat anticipation linked to fear circuit activation in humans and animals [47, 48]. Although alterations in arousal or reactivity are commonly reported in humans with PTSD [1], empirical evidence for exaggerated startle response is mixed [49, 50]. A meta-analysis that compared adults with and without PTSD indicated only modest increases in baseline startle reactivity in PTSD patients [51]. One potential reason for these moderate associations is that stress may differentially alter startle reactivity in subgroups (i.e., males versus females) of PTSD patients with distinct trauma-related pathology or trauma histories [52]. For example, after exposure to a terrorist attack, women reported higher levels of re-experiencing symptoms and exaggerated startle response than men [53]. In contrast, in female victims of childhood corporal punishment and partner aggression, higher PTSD symptom scores were related to lower startle reflex [28].

In this study, only male rats with a history of alcohol consumption showed stress-induced increases in startle reactivity after bobcat urine exposure. In agreement with these findings, previous studies have reported that individuals with comorbid PTSD and alcohol misuse exhibit more severe PTSD symptoms [54, 55]. Startle reactivity was tested 6 days after the last drinking session (i.e., not during intoxication). Furthermore, it is unlikely that animals in this study consumed quantities of alcohol sufficient to produce alcohol dependence or withdrawal. Contrary to prior reports [56], we found that male rats consumed more alcohol than females, and neither males nor females significantly escalated ethanol intake during the 5 weeks of intermittent 20% ethanol drinking. It is worth mentioning that our animals were pair-housed throughout the experiment, whereas in most studies using this protocol rats were housed individually [42, 57]. While the procedure used here precludes correlations between alcohol intake and startle reactivity in individual animals, it has the benefit of avoiding potential confound effects of social isolation.

Bobcat urine exposure reduced startle reactivity in females, regardless of alcohol drinking history. It is important to note that we did not measure baseline ASR to counterbalance the groups before the test session, but it is unlikely that random assignment of animals would select for rats with different startle reactivity. Although contrary to our initial hypothesis, prior work from others has reported lower startle reactivity in female rats after inescapable tail shock stress [58], an effect that was blocked by a systemic IL-1β injection in intact females, but not in ovariectomized females [59].

Our group previously reported that bobcat urine exposure increased anxiety-like behavior in male rats 2 days and 5 days later [46]. Here, we show that bobcat urine exposure did not alter anxiety-like behavior on day 21, i.e. 17 days post-stress, in male or female alcohol-naïve rats. Males with a history of alcohol drinking spent less time in the open arms of the elevated plus-maze and exhibited reduced general activity on the EPM relative to alcohol-drinking females. A similar reduction in general activity on the EPM was described in Long-Evans male rats after a 6-week intermittent-access to 20% ethanol in comparison to females exposed to the same procedure [56]. The higher open arm times in females may reflect sex differences in stress/fear coping strategies rather than “less anxiety-like behavior”. A previous study using factor analysis suggested that the behavior of female rats in the EPM is characterized more by activity than anxiety [60]. Also, females are four times more likely than males to display fear in the form of rapid movements instead of freezing in traditional models of Pavlovian fear conditioning [61].

Predator odor exposure increased corticosterone levels in females relative to unstressed controls and stressed males. Our group has previously shown that male Avoiders exhibit attenuated corticosterone response immediately following exposure to bobcat urine [46]. Differently from this study, the animals used here to evaluate endocrine responses were exposed to bobcat urine in a clean cage instead of in the 3-chamber apparatus, therefore they were not indexed as Avoiders or Non-Avoiders. Similar to our results, female rats exposed to acute restraint stress exhibit higher adrenocorticotropic hormone (ACTH) and corticosterone responses than males, in addition to significantly higher c-fos mRNA expression in the paraventricular nucleus of the hypothalamus (PVN) [62]. Conversely, TMT exposure elicits similar increase in circulating corticosterone in both male and female Wistar rats [63], indicating that distinct predator odors (i.e., natural olfactory stimuli or synthetic olfactory stimulus) can produce different responses. Future studies will determine whether higher corticosterone levels in females after bobcat urine exposure mediate stress-induced suppression of startle reactivity. Interestingly, suppression of ASR following combined cat exposure and saline injection in male rats can be blocked by substituting the injected saline with a glucocorticoid receptor antagonist [64].

## Conclusions

We report robust sex differences in behavioral and endocrine responses to bobcat urine exposure in adult Wistar rats. Males and females exposed to predator odor displayed blunted weight gain 24 hours post-stress, but only a subset of stressed animals exhibited avoidance behavior. Chronic moderate alcohol drinking increased traumatic stress reactivity in males but not females. Predator odor stress reduced startle reactivity in females relative to unstressed females and stressed males, regardless of alcohol drinking history. Furthermore, females exhibited higher increases in circulating corticosterone concentrations immediately following predator odor stress compared to males.

## Perspectives and significance

Sex differences in traumatic stress responses are among the most widely reported phenomena in epidemiological and clinical studies. To the best of our knowledge, the findings reported here are the first to provide evidence that a history of chronic moderate alcohol drinking differentially modulates predator odor stress reactivity in male and female rats. Our data support the notion that females rather respond differently to trauma and open doors for future work aimed at testing the neurobiology underlying sex differences in traumatic stress reactivity.

## Declarations

### Ethics approval

All procedures were approved by the Institutional Animal Care and Use Committee of the Louisiana State University Health Sciences Center and were in accordance with the National Institute of Health Guidelines.

## Consent for publication

Not applicable.

## Availability of data and materials

All data are available from the corresponding author upon request.

## Authors’ contributions

LA-S conceived and planned the study, performed the experiments, analyzed data and wrote the paper. CLS performed the experiments. NWG conceived and planned the study, supervised the project and contributed to the final version of the manuscript. All authors read and approved the final manuscript.

## Acknowledgments

The authors thank Curtis Vande Stouwe for his technical support.

This study was supported by the National Institutes of Health grants R01AA023305, R01AA026531, and the Cohen Veterans Bioscience Foundation. This work was also supported in part by Merit Review Award #I01 BX003451 (NWG) from the United States (U.S.) Department of Veterans Affairs, Biomedical Laboratory Research and Development Service.

